# Regulation of Ascorbic acid by gonadotropins in rat Ovary

**DOI:** 10.1101/665752

**Authors:** Shah Saddad Hussain, Taruna Arora, Madan Mohan Chaturvedi, Sharmila Basu-Modak, Rajesh Chaudhary, Kambadur Muralidhar

**Affiliations:** University of Alabama at Birmingham, Alabama, 35294-0006; Department of Zoology, University of Delhi, Delhi-110007; School of Life Sciences, University of Hyderabad, Hyderabad-500046; Department of Biomedical Science, Acharya Narendra Dev College, University of Delhi, Delhi-110007

**Keywords:** Gonadotropins, L-Gulonate dehydrogenase, ovarian metabolism

## Abstract

The dose dependent depletion of ovarian Ascorbic acid (AA) in rat ovaries, has been used as a bioassay for measurement of Luteinizing Hormone (LH). However, the mechanism of action of gonadotropin (LH, FSH) on ascorbic acid depletion is not completely clear in biochemical terms. To elucidate the mechanism, we looked for the pathways; one, where L-<u>Gu</u>lonate <u>D</u>e<u>h</u>ydrogenase (L-GuDH) catalyzes the conversion of L-Gulonic acid (L-GuA) to L-Xylulose, and, in the second the pathway conversion of L-GuA to AA, in a cats, dogs and Rats. Kinetic analysis of the enzyme L-GuDH *in vitro* showed the inhibitory effect of AA on L-GuDH. Therefore, we hypothesized that gonadotropins (FSH and LH) may regulate the L-GuDH maintain level of AA in ovary. LH administration to super-ovulated immature female rats caused depletion of ovarian AA but did not result in any change in the specific activity of the ovarian L-GuDH. Further, we administrated a surrogate FSH like hormone (PMSG) to immature female rats which, resulted in increased specific activity of ovarian L-GuDH. However, microarray data on RNA from ovaries exposed to FSH like hormone such as Pregnant Mare serum Gonadotropin (PMSG) did not reveal any increased expression of L-GuDH transcript. It is therefore concluded from the results obtained that; that neither LH, in decreasing the ovarian AA, nor FSH, in increasing the ovarian AA do so by regulating the activity of enzyme L-GuDH at transcriptional level. The results obtained have also been discussed by giving emphasis on the mechanism of ovarian ascorbic acid regulation of LH and FSH.

## Introduction

The gonadotropins, FSH (Follicle Stimulating Hormone) and LH (Luteinizing Hormone) secreted by anterior pituitary gland in response to intrinsic as well as extrinsic influences, stimulate a variety of ovarian functions including steroidogenesis, resumption of oocyte meiosis, ovulation, maintenance and formation of corpus luteum (Lindner *et al*. 1974). In males, the role of LH is mainly to control the synthesis and secretion of testosterone by Leydig cells. In females, it plays crucial role in ovulation and sustenance of the corpus luteum in the luteal phase of the female reproductive cycle (Dufau 1998). FSH is indispensable for both male and female gametogenesis. Pregnant Mare Serum Gonadotropins (PMSG) is a glycoprotein hormone secreted from endometrial cups of pregnant mare following 35 days of gestation (Papkoff 1974).PMSG exhibits both LH and FSH activities in heterologous animal species (Cole *et al*. 1940; Combarnous 1992) while being mostly LH like in equids (Urwin & Allen 1982). In the immature male rat, PMSG increases testicular weight, the enlargement of seminal vesicles (Cole *et al*. 1932; Raacke *et al*. 1957) The dual function of this placental gonadotropin has generated extensive interest with regard to mechanism of action *in vivo* (Licht *et al*. 1979; Moudgal & Papkoff 1982). The function of gonadotropins, (LH and FSH) and thyrotropin has been found to be mediated through the second messenger cyclic AMP (Moudgal *et al*. 1971). Gonadotropins have also been shown to elicit some non-steroidogenic responses from the ovaries like incorporation of Uridine to RNA, and incorporation as well as transport of proteins into ovary or increasing the poly A length of ovarian transcripts (Ahren, K, Hamberger, L, Rubinstein 1969; Jarlstedt *et al*. 1973; Selstam & Nilsson 1974; Prasad *et al*. 1978). One of oldest and the most widely used bioassays for LH measurement by LH induced depletion of ovarian ascorbic acid in super-ovulated immature rats, first observed by Parlow (Parlow 1961). Some factors other than LH also influence ascorbic acid level in ovaries (Miszkiel 1999). Vasopressin also possesses appreciable activity like that of LH (Mccann & Taleisnik 1960). Purified preparation of other anterior pituitary hormones failed to affect the levels of ascorbic acid in ovary (Mccann & Taleisnik 1960; Schmidt-Elmendorf & Loraine 1962). In animals like rat and mice it could be proposed that LH must be affecting, either biosynthesis or catabolism or both, of ascorbic acid in the ovary. Another possibility is that LH may be accelerating transfer of ascorbic acid from ovary to venous circulation. Ascorbic acid metabolism involves more than one metabolic pathway. In the biosynthetic pathway of ascorbic acid, L-gulonolactone get converted to L-ascorbic acid via 2-keto-L-gulonolactone and the latter reaction is catalyzed by L-gulonolactone oxidase (gulo)(Chatterjee *et al*. 1960). In the animals such as mice and rats that are capable of synthesizing AA, therefore, two pathways are regulated through the intermediate L-Gulonate either towards ascorbic acid formation or towards the Xylulose formation. The question that arises is what external factors control this channeling of L-Gulonate into either of the two possible pathways? Alternatively, one can envisage a condition where L-Gulonate is not transferred to ascorbate under normal condition. Under special condition, it is possible that L-GuDH is blocked, thus formation of xylitol is blocked and hence ascorbic acid concentration is increased. Does such a scenario happen in reverse in ovary? (Figure 1). How is the depletion of ovarian AA by LH regulated? Is there any role of L-GuDH in the regulation of AA depletion?. Vitamin C or ascorbic acid has been found to play an important role in reproductive process (Kramer *et al*. 1933). Parlow had shown the gonadotropin action of LH on ovarian ascorbic acid content in 1961, where he observed that there was marked decrease in ovarian ascorbic acid in the corpus luteum and possibly during lutenization itself (Parlow 1961). In order to explore the association between L-GuDH, ascorbic acid and hormonal action, we initiated the study by using rat model, in which we performed commonly used bioassay called Ovarian Ascorbic Acid Depletion Assay (OAAD) or Parlow bioassay.

**Figure 1:**
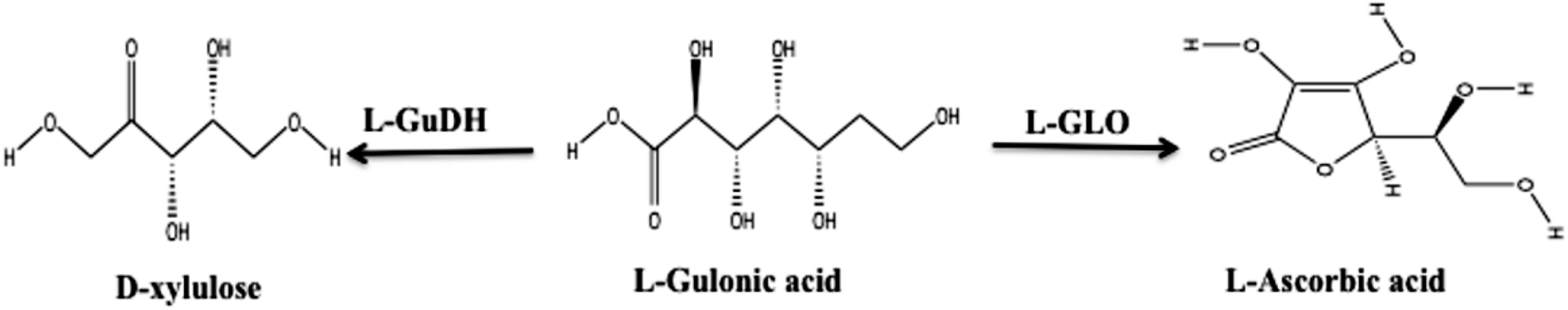
Metabolic fates of L-Gulonate. L-gulonate gets converted to L-ascorbic acid via 2-keto L-gulonolactone and this reaction is catalyzed by L-gulonolactone oxidase (gulo). L-gulonate gets converted to D Xylulose via L-GuDH.

## Materials and Methods

### Animals

Animal experiments were carried out according to the guidelines of CPCSEA,” Government of India, with Institutional Animal Ethics Committee IAEC approval DUZOO/IAEC-R/2011/17. Rats were of the Holtzman strain, housed in our animal house under controlled temperature (21–23°C) and with the light period adjusted to 12 h light and 12 h darkness. The animals had free access to standard pelleted laboratory animal food and tap water.

### Chemicals and Hormones

L-Gulonate-γ-lactone, NAD, PMSG, hCG, 2,6 dichloro phenol indophenol (monosodium salt)(DCPIP), theophylline and L-ascorbic acid were purchased from Sigma USA. All other chemicals used were of analytical grade

### Estimation of Protein

Protein estimation of the samples was performed at each step, according to the method given by Lowry(Lowry *et al*. 1951).

### Effect of hCG (LH like hormone) on ovaries

Super ovulated immature female rats of Holtzman strain 26-28 days old were used for the experiments. Rats were made super ovulated by injecting 50IU of PMSG (in 0.2ml of albumin phosphate buffer) subcutaneously twice 48 hours apart. 56 to 60 hours later the animals received 25IU of hCG in 0.2ml of albumin phosphate buffer again subcutaneously. On the 6^th^ day after the injection of hCG the animals were used for the experiment. On the day of experiment animals were divided into two groups control and experimental, 3 animals (n=3) in each group. The control animals received only albumin phosphate buffer and experimental animals received 60IU hCG in 0.2ml of albumin phosphate buffer intraperitoneally. Four hours later, the animals were taken for autopsy. Left ovaries were used for AA estimation and right ovaries for L-GuDH assay. Placing the ovaries on an ice-chilled Petri dish containing 5% metaphosphoric acid to stop the metabolic activities did subsequent processing for left ovary. The ovary was freed of surrounding adipose tissue and quickly weighed to the nearest mg precision balance. Slightly minced ovarian tissue was then homogenized in 3ml of 5% metaphosphoric acid–citrate solution, using a Potter-Elvehjem Teflon tissue grinder. The homogenate was then centrifuged at 10000 rpm for 30min at 4°c using a cold centrifuge (SIGMA). An aliquot of 3mL from the clear supernatant was used for the estimation of AA according as follows.

A standard calibration curve was constructed using 0-80 μg of ascorbic acid in 2 mL of 5% metaphosphoric acid–citrate solution. 2 mL of the dye solution DCPIP was delivered quickly into the tubes mixed well and absorbance was read at 540nm at 20^th^ ± 2 seconds after mixing in the spectrocolorimeter. Ovarian ascorbic acid was estimated essentially as per the assay of Parlow (Parlow 1961).

### Effect of PMSG (FSH like hormone) on ovaries

For the experiment, 25-26 day old immature female rats (Holtzman strain) were used. Two groups of animals were taken, 4 animals (n=4) in each group. The control animals received only albumin phosphate buffer subcutaneously while the experimental group of animals received a single injections of 50IU of PMSG. Two days later, animals were sacrificed. The left ovary was processed for the ascorbic acid estimation and RNA extraction while the right ovary was processed for the enzyme assay and protein estimation. After autopsy the right ovaries of the rats were immediately removed. Subsequent processing of right ovaries was done by placing the ovaries on an ice-chilled Petri dish having 50mM PB, pH 7.5 which contained 1mM PMSF (Extraction buffer). The ovary was freed of surrounding adipose tissue and quickly weighed to the nearest mg precision balance. Slightly minced ovarian tissue was then homogenized using a Potter-Elvehjem Teflon tissue grinder in 50mM PB, pH 7.5 containing 1mM PMSF. Finally the homogenate was centrifuged at 10000 rpm for 30 min at 4°c using a cold centrifuge (SIGMA make). An aliquot of 1mL from the clear supernatant was used for the assay of GuDH and protein estimation.

### Measurement of L-GuDH Activity

L-GuDH the dehydrogenase activities of each samples of the bioassay were assayed by measuring the rate of change in NADH absorbance at 340 nm as described above. The amount of NADH produced as a result of dehydrogenase activity was calculated from the slopes of the curves obtained by change in absorbance against time interval at 340 nm as dictated by the assay method. Considering 6.220 as the micro molar extinction coefficient of NADH quantitated the units of GuDH present in one mL.

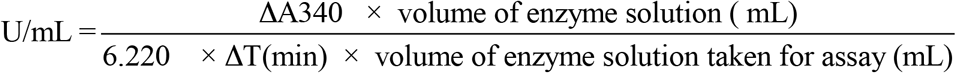

Specific activity or was calculated by dividing the value of U/mL by the concentration of protein.

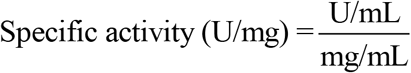

### Gene expression profiling following exposure to PMSG

Slides used: Agilent-Custom: Rat 8X60K designed by Genotypic Technology Private Limited (AMADID: 28279).

### RNA extraction and evaluation

RNA isolation was performed using Qiagen RNeasy Mini Kit as per the manufacturer’s instructions. The concentration and purity of the RNA extracted were evaluated using the Nano drop Spectrophotometer (Thermo Scientific; 1000). The integrity of the extracted RNA were analysed on the Bio analyzer (Agilent; 2100). RNA quality was based on the 260/280 values (Nano drop), rRNA 28S/18S ratios and RNA integrity number (RIN) (Bio analyzer).

### RNA Labelling, Amplification and Hybridization

The samples were labeled using Agilent Quick-Amp labeling kit (Part number: 5190-0424). 500ng of total RNA was reverse transcribed using oligo dT based method. mRNA was primed with oligo dT primer tagged to T7 promoter sequence and converted into double stranded cDNA using MMLV-RT reverse transcriptase. Further, in the same reaction cDNA was *in-vitro* transcribed to cRNA using T7 RNA polymerase enzyme. During cRNA synthesis, Cy3 labeled Cytosine nucleotide was incorporated into the newly synthesized strands. Labeled cRNA thus obtained was cleaned up using Qiagen RNeasy columns (Qiagen, Cat No: 74106). The concentration and amount of dye incorporated were determined using Nanodrop. 600ng of labeled cRNA were hybridized on the array (AMADID: 28279) using the Agilent Gene Expression Hybridization kit (Part Number 5190-0404) in Sure hybridization Chambers (Agilent) at 65°C for 16 hours. Hybridized slides were washed using wash buffers (Part No: 5188-5327; Agilent). The hybridized and washed microarray slides were then scanned on a G2505C scanner (Agilent Technologies).

### Feature Extraction

Data extraction from Images was done using Feature Extraction software Version 11.5.1.1 of Agilent.

### Microarray Data Analysis

Images were quantified using Feature Extraction Software (Version-11.5, Agilent). Feature extracted raw data was analyzed using Gene Spring GX Version 12.0 software from Agilent. Normalization of the data was done in Gene Spring GX using the 75th percentile shift (Percentile shift normalization is a global normalization, where the locations of all the spot intensities in an array are adjusted. This normalization takes each column in an experiment independently, and computes the *n^th^* percentile of the expression values for this array, across all spots (where n has a range from 0-100 and n=75 is the median). It subtracts this value from the expression value of each entity) and normalized to Specific control Samples. Significant genes up and down regulated in test samples with respect to control sample were identified. Statistical t-test p-value was calculated based on volcano Plot. Differentially regulated genes were clustered using hierarchical clustering based on Pearson coefficient correlation algorithm to identify significant gene expression patterns. Genes were also classified based on functional category and pathways using DAVID (Database for Annotation, Visualization and Integrated Discovery) database tool (Huang *et al*. 2009). The microarray data are deposited at the Gene Expression Omnibus public repository (http://www.ncbi.nlm.nih.gov/geo) with accession number GSE 68676.

### Results and Discussion

The ovarian ascorbic acid depletion assay devised by Parlow (1961) is one of the most potent and reliable bioassays for LH. The calibration curve for ascorbic acid estimation showed that the method works well in the range of 0 to 80μg of ascorbic acid. (Figure 1)

A significant depletion of ascorbic acid content was observed in experimental group as compared to control group (Table1). With respect to the control animal 60IU hCG administered animals had ascorbic acid content depleted up to 65.13%. One of the known actions of pituitary Lutropin (Luteinizing hormone) is to cause depletion of ovarian ascorbate in the pseudo-pregnant rat (Parlow 1961). However, the actual role of AA in LH induced ovulation is not clearly known. The depletion of ovarian ascorbate by lutropin has been utilized as a sensitive bioassay of lutropin. This is reported to occur in corpora lutea within minutes of lutropin injection and exhibits a characteristic time sequence(Goldstein & Sturgis 1961). Role of vitamin C has been observed in steroidogenesis and peptide hormone production (Luck *et al*. 1995).

**Table 1:** Ability of hCG to deplete ovarian ascorbic acid

| Group | Treatment | Ascorbic acid ($\mu\text{g}/100\text{gm}$ body wt) | % age Depletion of AA |
| --- | --- | --- | --- |
| Control (n=3) | Only buffer | 158.36±14.89 | 0 |
| Experimental (n=3) | 60 IU hCG | 55.215±8.419 | 65.13 |

$p < 0.005$ ; $df = 6$ , Mean ( ± SE; $n = 3$ )

When the enzyme (L-GuDH) was assayed, it was observed that the rate of change in absorbance of NADH with respect to time at 340nm remained almost same in control and experimental groups in both the trials (Figure 2). NADH represents the activity of GuDH. Further, the difference in Nano moles of NADH formed per minute per 100mg of ovarian weight or per mg protein, in experimental and control groups, was not significant (data not shown). Hence it can be concluded that LH does not regulate the activity of L-GuDH in the ovary. It was also observed that this effect of LH was Cyclohexiimide insensitive (Arora *et al*. 2012). Therefore induced or constitutive protein synthesis is not required for this depleting effect of LH on ovarian ascorbic acid. Earlier work had shown that the administration of LH to Parlow rats results in increase in ovarian content poly A rich RNA (Muralidhar *et al*. 1976; Prasad *et al*. 1978) and that this effect could be observed within 30 minutes of LH administration. Hence the effect of LH on rat ovary appears to be both metabolic (non-genetic) and developmental (genetic) depending on the animal model.

**Figure 2:**
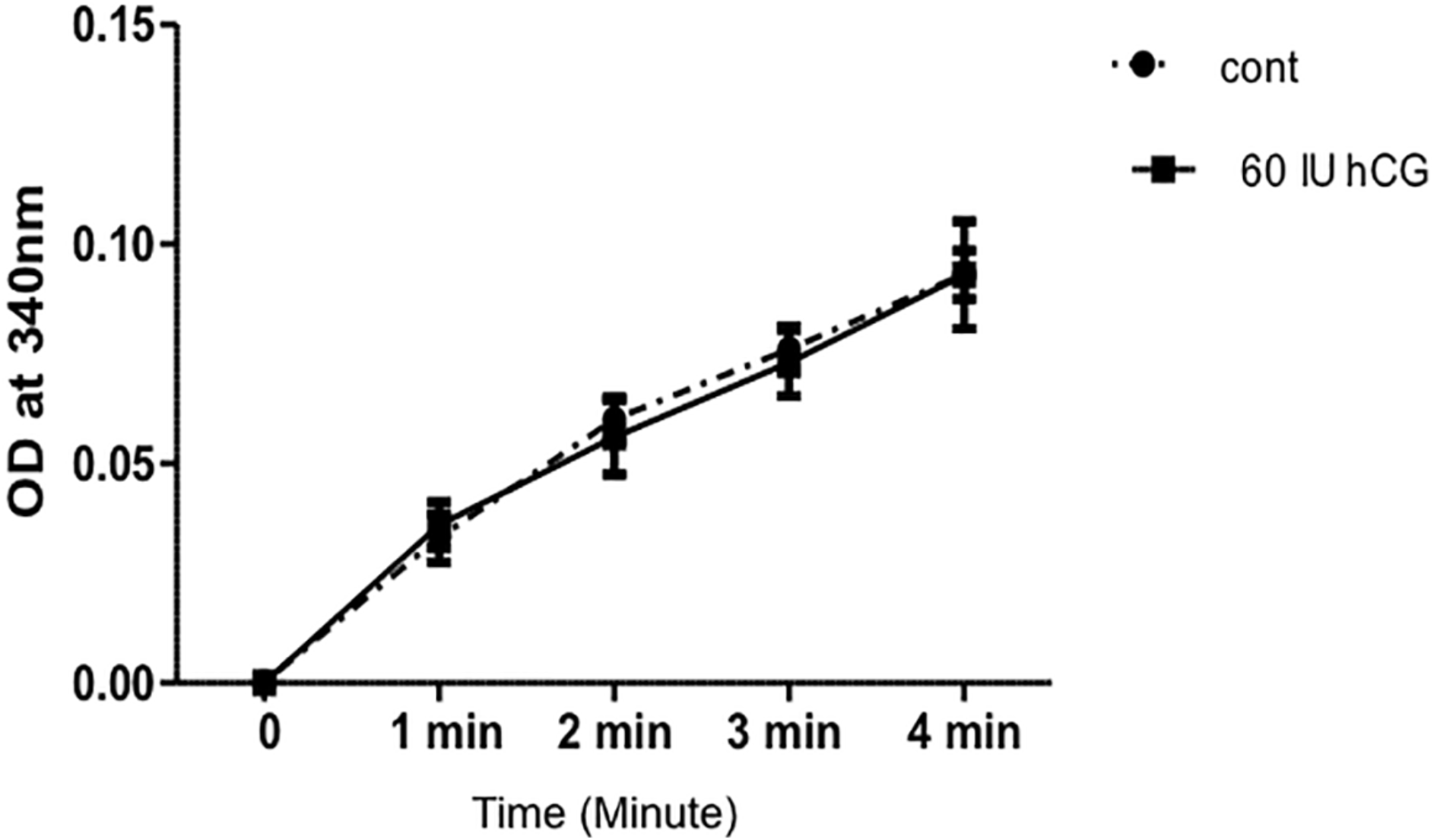
L-GuDH activity curve showing change in NADH absorbance at wavelength 340 nm in control and 60 IU hCG treated group when equal amount of ovarian extract was taken for assay.

In the experiment on effect of PMSG on rat ovarian ascorbic acid content, however, results indicate that administration of PMSG (a long acting FSH-like hormone) resulted in increase in gonado-somatic index (Figure 3A) and increase in ascorbic acid content (Figure 3B). Further, the same treatment also resulted in increase in enzyme activity (Figure 4A), total protein content (Figure 4B) and in specific activity (Figure 4C). It is known that during follicular growth ovulation and formation of corpora lutea ECM undergoes constant changes, so it requires high level of collagen and ascorbic acid fulfills this demand by promotion of biosynthesis of collagen (Kramer *et al*. 1933). The elevation in ascorbic acid levels can be explained by this mechanism. Growing corpus luteum requirement of collagen is fulfilled by ascorbate as it serves as a cofactor in collagen synthesis (Luck *et al*. 1995). So ascorbic acid is playing direct role in the function of corpus luteum. (Byrd *et al*. 1993).

**Figure 3:**
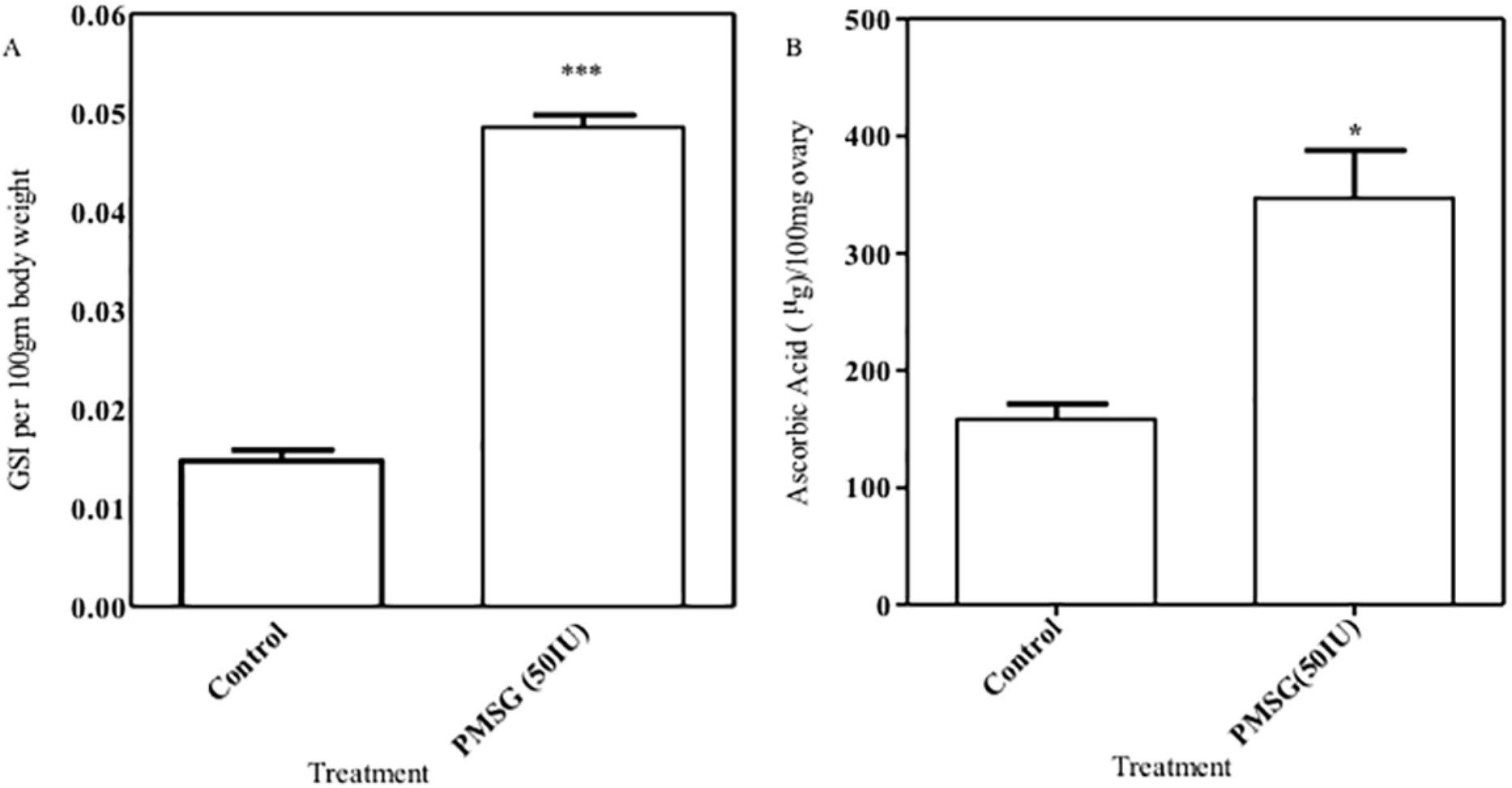
Effect of PMSG on ovary. A) GSI (Gonado Somatic Index) per 100gm body weight 48 hr after 50 IU of PMSG injections. B) The effect of PMSG on ovarian ascorbic acid content in immature rat.

**Figure 4:**
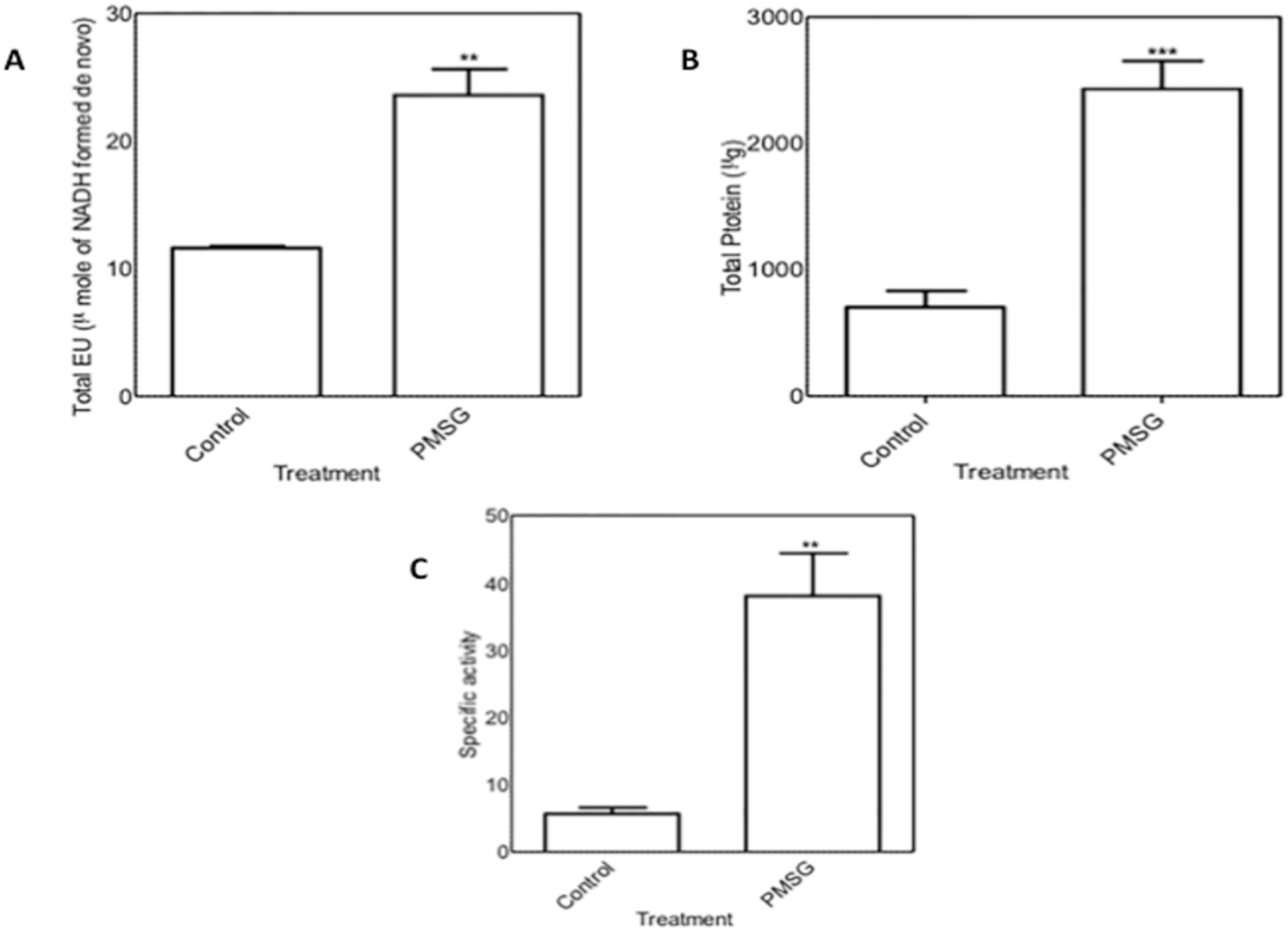
Effect of 50 IU PMSG on ovary A) Total L-GuDH units (μ moles of NADH formed) of ovary after PMSG (50IU) injection. B) Total Protein content of ovary 48 hrs after injection of PMSG (50IU). C) Specific activity of L-GuDH in ovary after PMSG (50IU) injection.

Ascorbic acid inhibits the L-Gulonate dehydrogenase in *in vitro* condition (Sharma *et al*. 2013). It was interpreted that the high levels of ascorbic acid inhibited the enzyme in vivo. However molecular docking studies also indicate that ascorbic acid can bind GuDH (unpublished work). It was also observed that there was no increase in specific activity of ovarian L-Gulonic acid dehydrogenase (L-GuDH) activity during the decrease in ascorbic acid under *in vivo* conditions. In the case of FSH action, however, simultaneous increase in both ascorbate levels and L-Gulonate dehydrogenase activity (Figure 3 and 4) was observed.

To understand the regulatory mechanism of FSH and LH on Ascorbic acid and L-GuDH, it is imperative to explore different pathways and the genes involved in ovarian functions. Hence a microarray analysis was undertaken on ovaries exposed to PMSG *in vivo*. The quality of RNA preparation was assessed and found to be good for further study.

The global expression profile of rat ovaries challenged with PMSG was compared with controls (without PMSG). Probe sets whose intensities were normalized and filtered, then were subjected to identifying significantly differentially expressed genes using t-test analysis by comparing the log2 (normalized signal) of two different samples of ovary, and 1904 transcripts were identified to be differentially up regulated at the p ≤ 0.05 level. Taking a FC ≥ 1.5 or ≤ 0.6 and the p ≤ 0.05 significance level as the criteria, 1904 transcripts showed differentially up regulated and 1414 showed down regulated expression.

Gene Ontology (GO) and Kyoto Encyclopedia of Genes and Genomes (KEGG) pathway analyses of all annotated DE gene lists were carried out by using the Database for Annotation, Visualization and Integrated Discovery (DAVID). GO annotation mapping revealed that the genes assigned GO terms for reproductive cellular process, biological regulation, regulation of MAPK cascade; somatic stem cell division; organ development; embryonic morphogenesis; reproductive process in a multicellular organism; regulation of immune response; negative regulation of immune response; regulation of developmental process; regulation of cellular process; regulation of cell division; regulation of cell cycle; biological regulation; regulation of response to stress, developmental process, cell communication and signal transduction and so on (Table 2). DE transcripts were involved in some important pathways. Including the estrogen-signaling pathway, MAPK signaling pathway and ovarian steroidogenesis, and ascorbate and aldrate metabolism (figure 5)

**Figure 5:**
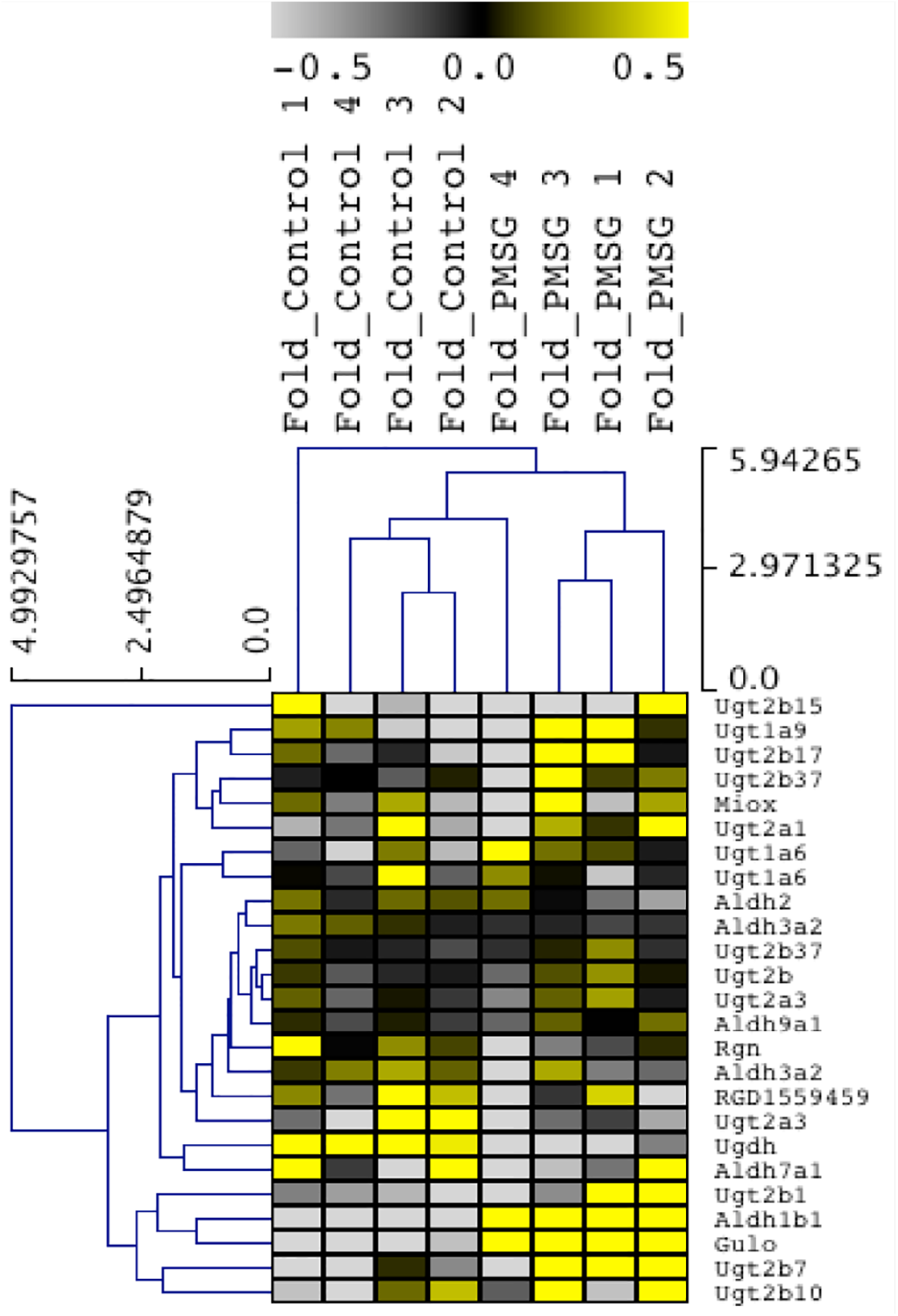
Heat map-showing changes in expression of genes related to different pathways. A) Ovarian steroidogenesis. B) Ascorbate aldrate pathway. C) Estrogen signaling Pathway. D) MAPK signaling Pathway. Heat maps were constructed using average of raw signals for each gene in microarray data. Expression fold values are provided in terms of log base 2. Differential expression calculated using p ≤ 0.05 levels and taking a FC ≥ 1.5 or ≤ 0.6 and the p ≤ 0.05 significance level as the criteria.

**Table 2:** Functional category enrichment analysis based on Gene Ontology terms.

| Biological Process | GO Term ID | Counts | % of Differentially Regulated genes | P Value |
| --- | --- | --- | --- | --- |
| Developmental process | GO:0032502 | 247 | 18.69795609 | 0.00 |
| Biological regulation | GO:0065007 | 404 | 30.58289175 | 0.00 |
| Organ development | GO:0048513 | 150 | 11.35503407 | 0.00 |
| Regulation of cellular process | GO:0050794 | 332 | 25.1324754 | 0.00 |
| Reproduction | GO:0000003 | 60 | 4.542013626 | 0.00 |
| Sexual reproduction | GO:0019953 | 37 | 2.800908403 | 0.00 |
| Reproductive process | GO:0022414 | 59 | 4.466313399 | 0.00 |
| Multicellular organism reproduction | GO:0032504 | 41 | 3.103709311 | 0.01 |
| Reproductive process in a multicellular organism | GO:0048609 | 41 | 3.103709311 | 0.01 |
| Regulation of response to stress | GO:0080134 | 26 | 1.968205905 | 0.01 |
| Reproductive structure development | GO:0048608 | 18 | 1.362604088 | 0.01 |
| Reproductive developmental process | GO:0003006 | 28 | 2.119606359 | 0.01 |
| Embryonic morphogenesis | GO:0048598 | 30 | 2.271006813 | 0.02 |
| Regulation of developmental process | GO:0050793 | 52 | 3.936411809 | 0.09 |
| Reproductive cellular process | GO:0048610 | 16 | 1.211203634 | 0.12 |
| Regulation of MAPKKK cascade | GO:0043408 | 11 | 0.832702498 | 0.18 |
| Negative regulation of immune response | GO:0050777 | 4 | 0.302800908 | 0.33 |
| Somatic stem cell division | GO:0048103 | 2 | 0.151400454 | 0.36 |
| Regulation of cell division | GO:0051302 | 4 | 0.302800908 | 0.41 |
| Regulation of cell cycle | GO:0051726 | 14 | 1.059803179 | 0.75 |
| Two-component signal transduction system (phosphorelay) | GO:0000160 | 1 | 0.075700227 | 1.00 |

### Differential Gene Expression Analysis

The functions and pathway analysis was done for the differentially regulated genes using DAVID database. Pairwise analysis of overall gene expression profiles between the 8 samples of immature female rat ovary of 26 to 28 days old rat was done. We found that 1904 genes were up regulated more than 1.5 fold and 1414 genes were found to be down regulated below 1.5 fold. Based on gene ontology study we have chosen some of the functionally important genes in oogenesis. It can be noticed that only the lactonase gene (*Gulo*) expression increased but not that of gene for L-GuDH (Figure 6). No increase in gene expression levels of the gene for L-GuDH was observed. From current observations, it appears that actions of PMSG (FSH like) on specific activity of L-Gulonate-3-dehydrogenase and ascorbic acid content are independent phenomena and L-Gulonate-3-dehydrogenase may not have a role in the regulation of ovarian ascorbic acid content. Stabilization of mRNA for the GuDH and/or post-translational modification/activation of the GuDH could also have a played a role in total increase in the GuDH specific activity. Indeed there are reports that an mRNA population not involved in translation could be reactivated by cytoplasmic poly A polymerase (Weill *et al*. 2012). There is also a possibility that in the case of ovaries exposed to PMSG, without increase in expression level of L-GuDH gene, the dormant mRNA got activated and hence we observed the increase in specific activity of the enzyme.? In fact bioinformatics studies have revealed potential phosphorylation sites in L-GuDH protein structure (Shah Saddad Hussain-unpublished results). More research work needs to be done to throw light on this problem. Hence it can be tentatively concluded that LH does not regulate this enzyme transcriptionally and FSH probably regulates this enzyme through post-transcriptional mechanisms.

**Figure 6:**
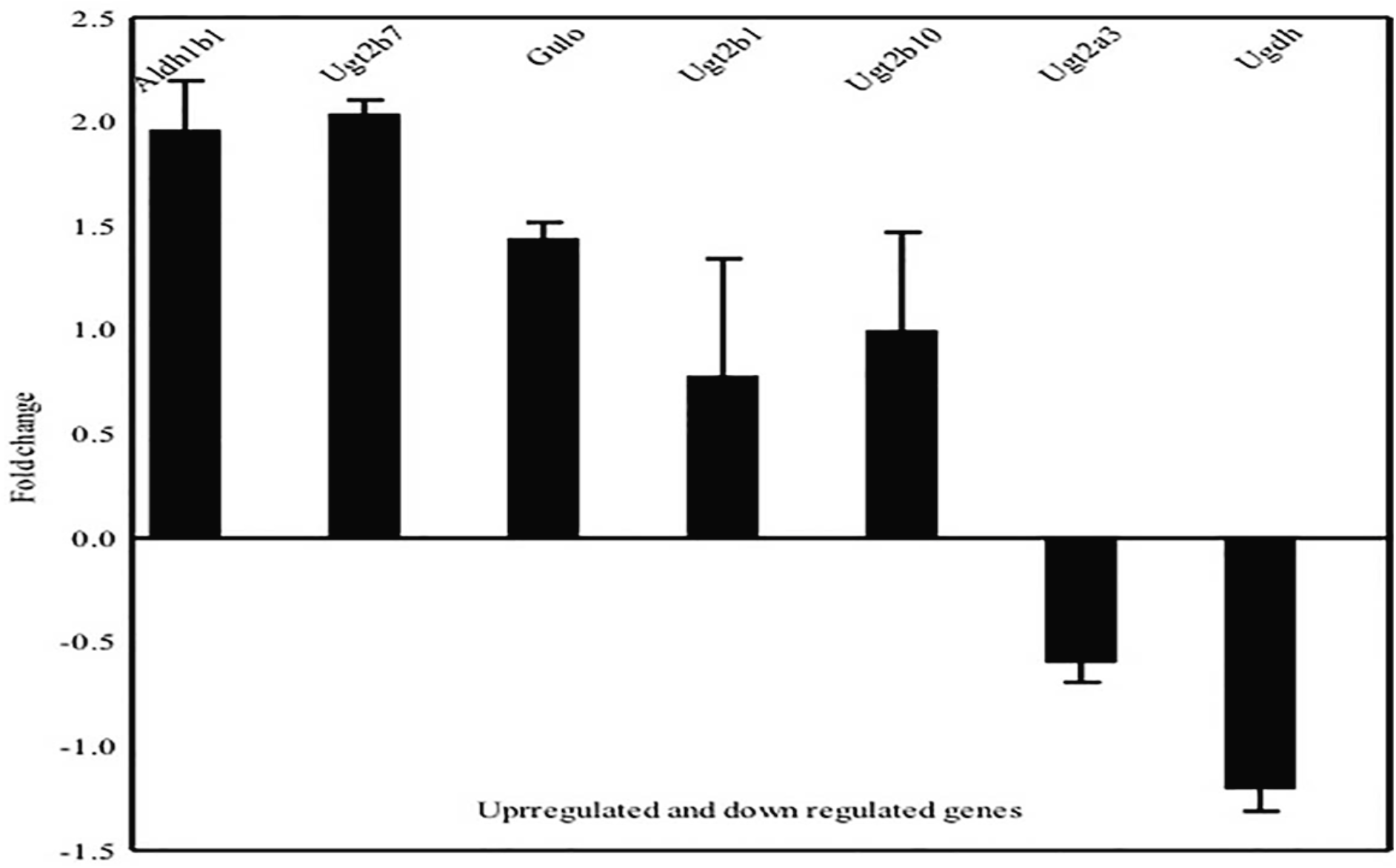
Functional pathway analysis of Micro array data. List of Genes up-regulated and down-regulated in Ascorbate metabolism pathway. Expression fold values are provided in terms of log base 2. Differential expression calculated using p ≤ 0.05 levels and taking a FC ≥ 1.5 or ≤ 0.6 and the p ≤ 0.05 significance level as the criteria.

## Disclosure statement

The authors have nothing to disclose.

## Funding

J.C Bose Fellowship to KM from DST, Government of India and funds from CSIR, Government of India to KM, SBM and SH during the project period.

## Author contributions

SSH and KM conceptualized and conceived the project; SSH and KM designed the studies and interpreted the result; SSH and KM wrote the manuscript; SSH executed the project; TA performed experiment; MMC Provided experimental facility and supervised, MMC SBM and RC reviewed and edited manuscript.

## Acknowledgements

Work reported here was supported by funds from the JC Bose fellowship grant to KM by DST (Govt of India) and CSIR funds in the form of EMR project to SBM in which SSH was a SRF. Financial help is hence gratefully acknowledged.

## Notes

#### Summary of Updates

Name of one of author : Sharmila B Modak to Sharmila Basu-Modak

https://www.ncbi.nlm.nih.gov/geo/query/acc.cgi?acc=GSE68676

## References

Ahren, K, Hamberger, L, Rubinstein L 1969 Actue in vivo and in vitro effects of gonadotrophins on the metabolism of the rat ovary. In The Gonads, K.W. McKer, p 327.

Arora T, Nath R, Vashishtha N, Kumari N, Hussain S & Muralidhar K 2012 Mechanistic investigations of the dual activity of Gonadotropins using target tissue cAMP level as a response parameter. World J Life Sci. and Medical Research 2 11–19.

Byrd JA, Pardue SL & Hargis BM 1993 Effect of ascorbate on luteinizing hormone stimulated progesterone biosynthesis in chicken granulosa cells in vitro. Comparative Biochemistry and Physiology. Comparative Physiology 104 279–281.

Chatterjee IB, Chatterjee GC, Ghosh NC, Ghosh JJ & Guha BC 1960 Biological synthesis of L-ascorbic acid in animal tissues: conversion of L-gulonolactone into L-ascorbic acid. The Biochemical Journal 74 193–203.

Cole HH, Guilbert HR & Goss H 1932 Further considerations of the properties of the gonad stimulating principle of mare serum. American Journal of Physiology -- Legacy Content 102 227–240.

Cole HH, Pencharz RI & Goss H 1940 On The Biological properties of highly purified gonadotropin from Pregnant Mare Serum. Endocrinology 27 548–553. (doi:10.1210/endo-27-4-548)

Combarnous Y 1992 Molecular basis of the specificity of binding of glycoprotein hormones to their receptors. Endocrine Reviews 13 670–691. (doi:10.1210/edrv-13-4-670)

Dufau ML 1998 The luteinizing hormone receptor. Annual Review of Physiology 60 461–496. (doi:10.1146/annurev.physiol.60.1.461)

Goldstein DP & Sturgis SH 1961 Luteinizing hormone-induced depletion of ascorbic acid in the rat ovary. The American Journal of Physiology 201 1053–1056.

Huang DW, Sherman BT & Lempicki RA 2009 Systematic and integrative analysis of large gene lists using DAVID bioinformatics resources. Nature Protocols 4 44–57. (doi:10.1038/nprot.2008.211)

Jarlstedt J, Nilsson L, Hamberger L & Ahren K 1973 Effects of gonadotrophins and cyclic 3’,5’-AMP on in vitro incorporation of (3 H)uridine into RNA of the prepubertal rat ovary. Acta Endocrinologica 72 771–785.

Kramer MM, Harman MT & Brill AK 1933 Disturbances of reproduction and ovarian changes in the guinea-pig in relation to vitamin C deficiency. American Journal of Physiology -- Legacy Content 106 611–622.

Licht P, Gallo AB, Aggarwal BB, Farmer SW, Castelino JB & Papkoff H 1979 Biological and binding activities of equine pituitary gonadotrophins and pregnant mare serum gonadotrophin. The Journal of Endocrinology 83 311–322.

Lindner HR, Tsafriri A, Lieberman ME, Zor U, Koch Y, Bauminger S & Barnea A 1974 Gonadotropin action on cultured Graafian follicles: induction of maturation division of the mammalian oocyte and differentiation of the luteal cell. Recent Progress in Hormone Research 30 79–138.

Lowry OH, Rosebrough NJ, Farr AL & Randall RJ 1951 Protein measurement with the Folin phenol reagent. The Journal of Biological Chemistry 193 265–275.

Luck MR, Jeyaseelan I & Scholes RA 1995 Ascorbic acid and fertility. Biology of Reproduction 52 262–266.

Mccann SM & Taleisnik S 1960 Effect of luteinizing hormone and vasopressin on ovarian ascorbic acid. The American Journal of Physiology 199 847–850.

Miszkiel G 1999 Original article Concentrations of catecholamines, ascorbic acid, progesterone and oxytocin in the corpora lutea of cyclic and pregnant cattle.

Moudgal NR & Papkoff H 1982 Equine luteinizing hormone possesses follicle-stimulating hormone activity in hypophysectomized female rats. Biology of Reproduction 26 935–942.

Moudgal NR, Moyle WR & Greep RO 1971 Specific binding of luteinizing hormone to Leydig tumor cells. The Journal of Biological Chemistry 246 4983–4986.

Muralidhar K, Prasad M, Adiga P & Moudgal N 1976 Stimulation of ovarian poly A rich RNA synthesis by human chorionic gonadotropin in vivo. IRCS Medical Science 4 88.

Papkoff H 1974 Chemical and biological properties of the subunits of pregnant mare serum gonadotropin. Biochemical and Biophysical Research Communications 58 397–404.

Parlow AF 1961 Human Pituitary Gonadotropins. CC Thomas, Spring field.

Prasad MSK, Muralidhar K, Moudgal NR & Adiga PR 1978 Effect of human chorionic gonadotrophin and ovine luteinizing hormone on rat ovarian macromolecular metabolism. The Journal of Endocrinology 76 283–292.

Raacke ID, Lostroh AJ, Boda JM & Li CH 1957 Some aspects of the characterization of pregnant mare serum gonadotrophin. Acta Endocrinologica 26 377–387.

Schmidt-Elmendorf H & Loraine JA 1962 Some observations on the ovarian ascorbic acid depletion method as a test for luteinizing hormone activity. The Journal of Endocrinology 23 413–421.

Selstam G & Nilsson L 1974 Effect of human menopausal gonadotrophin on amino acid transport in the prepubertal rat ovary. Acta Physiologica Scandinavica 90 798–800. (doi:10.1111/j.1748-1716.1974.tb05650.x)

Sharma V, Hussain SS, Pooja, Sonia, Nirmala K & Muralidhar K 2013 Buffalo Kidney L-Gulonate Dehydrogenase: Isolation and Kinetic Characterization. World J. Life Sc Med Res 3 8–14.

Urwin VE & Allen WR 1982 Pituitary and chorionic gonadotrophic control of ovarian function during early pregnancy in equids. Journal of Reproduction and Fertility. Supplement 32 371–381.

Weill L, Belloc E, Bava F-A & Mendez R 2012 Translational control by changes in poly(A) tail length: recycling mRNAs. Nature Structural & Molecular Biology 19 577–585. (doi:10.1038/nsmb.2311)

